# Comparative Genome Analysis Reveals Important Genetic Factors Associated with Probiotic Property in *Enterococcus faecium* strains

**DOI:** 10.1101/295881

**Authors:** Vikas C. Ghattargi, Meghana A. Gaikwad, Bharati S. Meti, Yogesh S. Nimonkar, Kunal Dixit, Om Prakash, Yogesh S. Shouche, Shrikant S. Pawar, Dhiraj Dhotre

**Author notes:** Corresponding Author(s) Dr. Dhiraj Dhotre, National Centre for Microbial Resource (NCMR), National Centre for Cell Science (NCCS), Pune- 411021, Maharashtra, India. Dr Shrikant S. Pawar, National Centre for Microbial Resource (NCMR), National Centre for Cell Science (NCCS), Pune- 411021, Maharashtra, India.

## Abstract

*Enterococcus faecium* though commensals in human gut, few strains provide beneficial effect to humans as probiotics, few are responsible for nosocomial infection and few as non-pathogens. Comparative genomics of *E. faecium* will help to reveal the genomic differences responsible for the said properties. In this study, we compared *E. faecium* strain 17OM39 with a marketed probiotic, non-pathogenic non-probiotic (NPNP) and pathogenic strains. The core genome analysis revealed, 17OM39 was closely related with marketed probiotic strain T110. Strain 17OM39 was found to be devoid of known vancomycin, tetracycline resistance genes and functional virulence genes. Moreover, 17OM39 is „less open‟ due to absence of frequently found transposable elements. Genes imparting beneficial functional properties were observed to be present in marketed probiotic T110 and 17OM39 strains. Additional, genes associated with colonization within gastrointestinal tract were detected across all the strains. Beyond shared genetic features; this study particularly identified genes that are responsible to impart probiotic, non-pathogenic and pathogenic features to the strains of *E. faecium.* The study also provides insights into the acquired and intrinsic drug resistance genes, which will be helpful for better understanding of the physiology of antibiotic resistance in *E. faecium* strains. In addition, we could identify genes contributing to the intrinsic ability of 17OM39 *E. faecium* isolate to be a potential probiotic.

The study has comprehensively characterized genome sequence of each strain to find the genetic variation and understand effects of these on functionality, phenotypic complexity. Further the evolutionary relationship of species along with adaptation strategies have been including in this study.

## BACKGROUND

The genus *Enterococcus* is one of the diverse and ecologically significant group and members of this genus are ubiquitously distributed in nature viz. animals, human gastrointestinal tract (GIT) and plants (Lam et al. 2012; Qin et al. 2012; Geldart and Kaznessis 2017; McKenney et al. 2016; dos Santos et al. 2015; Byappanahalli et al. 2012; Rasouli Pirouzian et al. 2012). *Enterococcus* plays an important role in the ripening of cheese products by lipolysis and proteolytic properties leading to the development of aroma and flavour (Rasouli Pirouzian et al. 2012). In Mediterranean region, *Enterococcus* spp have been used in the preparation of various meat and fermented milk products for centuries (dos Santos et al. 2015). Further, they also exhibit the beneficial property of bacteriocin production (Rasouli Pirouzian et al. 2012; dos Santos et al. 2015) presenting activity against potential pathogens viz. group D streptococci and *Listeria* in various foods and in GIT (Rasouli Pirouzian et al. 2012).

*E. faecium* is widely and extensively studied for its leading cause of nosocomial infections in humans (Guggenbichler et al. 2011). It is a gut commensal and acts as opportunistic pathogen due to a variety of virulence factors, including lipopolysaccharides and biofilm formation (Natarajan and Parani 2015). Their pathogenic nature is evident in urinary tract infections, endocarditis, and surgical wound infection, displaying its capability of causing a wide range of infections (Kajihara et al. 2015). Another remarkable character of *E. faecium* is its tolerance to many antimicrobial drugs (Coque et al. 2005; O‟Driscoll and Crank 2015). It has also acquired antibiotic-resistance gene against vancomycin and a multidrug resistance beta-lactamase gene (Miller et al. 2014). Besides, it has been shown that *E. faecium* is capable to acquire resistance to antibiotics by sporadic mutations and infections caused by these are normally difficult to treat (Hollenbeck and Rice 2012). The strains like Aus0004 and V583 are reported as pathogens (Lam et al. 2012).

Numerous studies in the last decade have validated the safety claim of *Enterococci* in foods and as probiotics (Huys et al. 2013; Araújo and Ferreira 2013; Giraffa 2002; Franz et al. 2003). The application of *Enterococci* as a starter culture e.g *Enterococcus faecium* SF68 (Switzerland) and as probiotic e.g *E. faecium* T110 (Japan) has been used widely (Benyacoub et al. 2005; Jong-Hoon Lee, Donghun Shin, Bitnara Lee, Hyundong Lee, Inhyung Lee 2017). Additionally, *E. faecium* T110 is a content of many commercial available probiotics and no cause of illness or death has been reported (Natarajan and Parani 2015). *E. faecium* is among one of the directly fed microorganism recognized by the Association of American Feed Control, 2016. It is permitted as probiotic supplement in the diet for poultry, dogs, piglets and mice (Kačániová et al. 2006; Kreuzer et al. 2012; Vahjen and Männer 2003; Benyacoub et al. 2005). Few strains of *E. faecium* (NRRL B-2354) act as surrogate microorganism used in place of pathogens for validation of thermal processing technologies (Kopit et al. 2014) and some are widely used as laboratory strains e.g. *E. faecium* 64/3 (Bender et al. 2015). These two strains are non-pathogenic and are used routinely without any known disease outbreak (VanRenterghem 2012).

Thus the diversity and plasticity of *E. faecium* are accountable for both probiotic and pathogenic nature (Abeijón et al. 2006; Hassanzadazar et al. 2014; Satish Kumar et al. 2011). The work described here elucidates the genetic divergence between strain 17OM39 with marketed probiotic, non-pathogenic, and pathogenic strains to identify the genes reported for pathogenicity, antibiotic resistance, and probiotic properties. An attempt has also been made to identify the genes present exclusively in probiotic strains.

## RESULTS AND DISCUSSION

### Strain selection

Whole genome sequences were downloaded from NCBI genome database and the strains were grouped into probiotic, non-pathogenic non-probiotic (NPNP) and pathogenic based on the literature survey (Table1). The pathogenic group had six strains: DO, Aus0004, Aus0085, 6E6, E39 and ATCC 700221(Lam et al. 2012; Qin et al. 2012; Geldart and Kaznessis 2017; McKenney et al. 2016). The first four were isolated from human blood and latter two from human stool. The non-pathogenic group had two strains: NRRL B-2354 and 64/3(Kopit et al. 2014; Bender et al. 2015). The probiotic group had the marketed strain T110 (Natarajan and Parani 2015) and strain 17OM39 that was isolated from healthy human gut (Ghattargi et al. 2018).

**Table 1.**
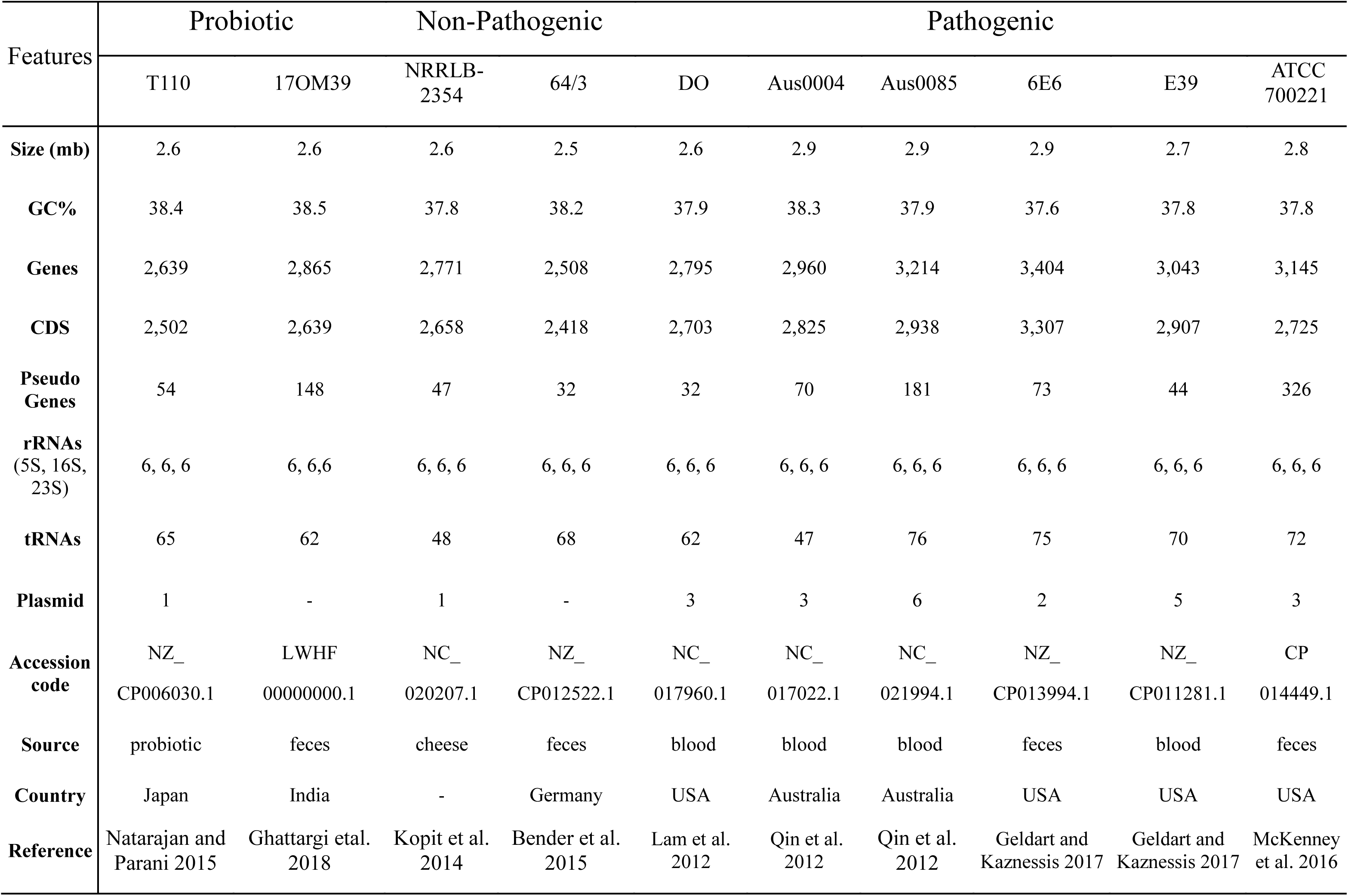
General Genome Features

### General genomic features

Genome sizes ranged from approximately 2.57–2.99 Mb with strain DO exhibiting the smallest and 6E6 the largest genome. Average GC content varied between 37.90 ± 0.65% and the strains with high G+C% do not have higher CDS, this contradicts with the results stated earlier (Bonacina et al. 2017). The genomic features of the strains under study are provided in Table 1. No significant differences (P ≤ 0.05, Kruskal–Wallis statistical test) could be noted between the groups with respect to their genome size, GC content, average number of genes and coding DNA sequence (CDS).

The RAST annotation has facilitated to determine the features, assigned to subsystems that are present in all organisms (Figure S1). The average number of annotated protein-encoding genes for the probiotic group was 2,570; for NPNP group was 2,639 and 3,093 for the pathogenic group. Annotation based on RAST for the strains under study suggests an abundance of carbohydrates and protein metabolism subsystems. The enriched carbohydrates metabolism is in agreement with the *E. faecium* ability to utilize a wide range of mono-, di-, oligo-saccharides (Devriese et al. 1987; Manero and Blanch 1999).

### Comparisons of 17OM39 with other E. faecium strains

The availability of complete *E. faecium* genomes has helped to define core, accessory and unique genomic features for all the strains. The comparison of strain 17OM39 with other strains of probiotic, NPNP and pathogenic strains, revealed 1935 (85.53 %) core genes, 526 (20.64%) accessory and 87(3.41%) unique genes. The numbers of shared genes are plotted as the function of number of strains (Figure 1). The figure shows that pan-genome size grew continuously with the addition of strains indicating an open pan-genome and the results are in accordance with previous study for *E. faecium* genome (Mikalsen et al. 2015). In contrast to the pan-genome, the size of the core-genome gradually stabilized (Mikalsen et al. 2015).

**Figure 1.**
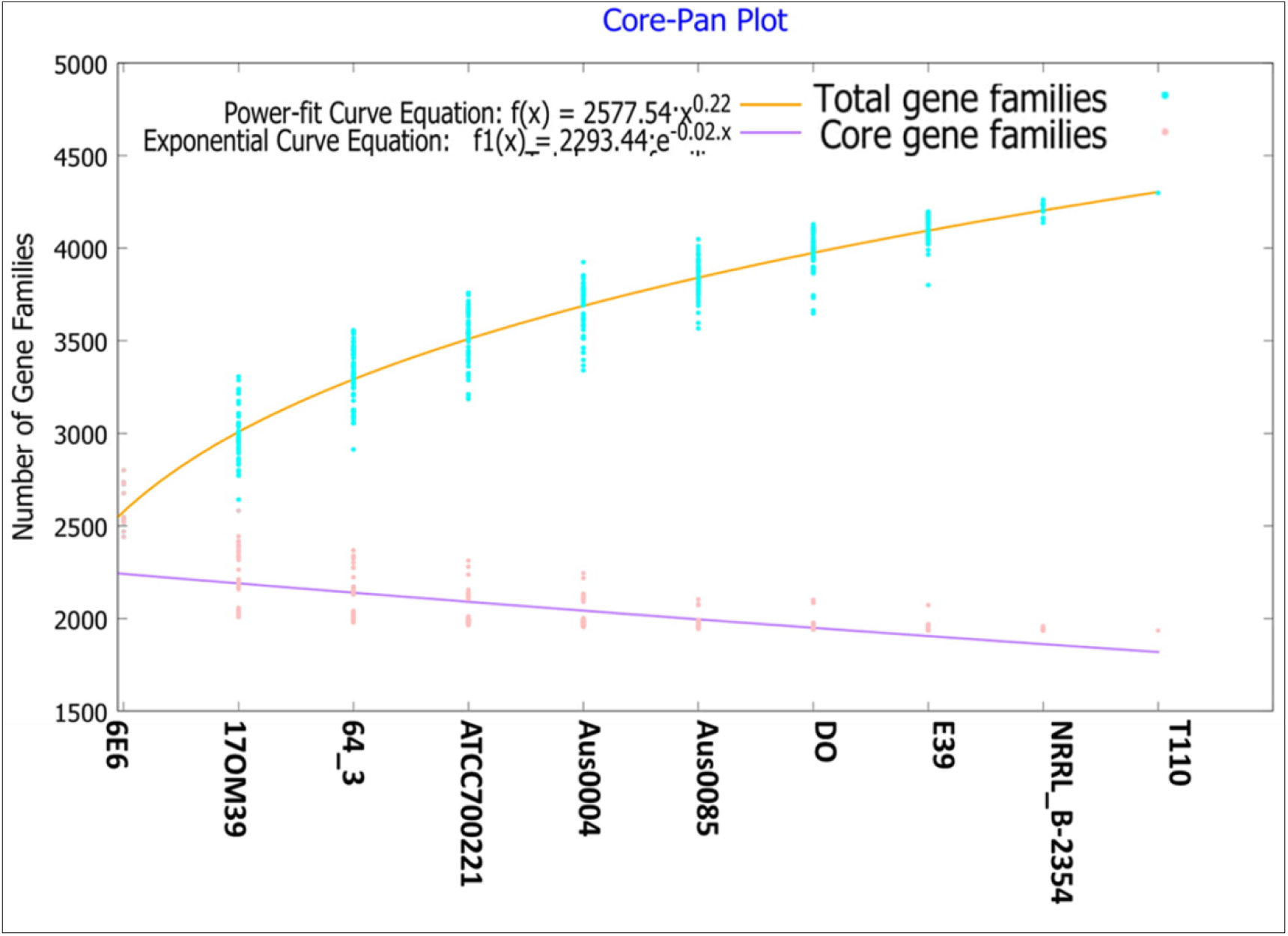
Core and pan genome for *E. faecium* strains. The number of shared genes is plotted as the function of number of strains (n) added sequentially. 1935 genes are shared by all 10 genomes. The orange line represents the least-squares fit to the power law function f(x)=a.x^b where a= 2577.54, b= 0.222602. The red line represents the least-squares fit to the exponential decay function f1(x)=c.e^(d.x) where c= 2293.44, d= -0.0232013.

### Pan-Genome Analysis

The pan genome analysis revealed the presence of 1935 core genes and 5718 accessory genes (Figure 2). The number of strain-specific genes observed was 67, 87, 10, 64, 62, 16, 13, 36, 54 and 14 for strains 17OM39, T110, NRRL B-2354, 64/3, DO, AUS0004, AUS0085, 6E6, E39 and ATCC700221, respectively (Figure 2). Identification of core, accessory and unique gene families by orthoMCL analysis revealed the proportion of known, hypothetical and uncharacterized proteins in these groups (Figure S2A). A large percentage (61.19%) of unique genes was assigned to an uncharacterized group and further studies are required to examine these unexplored attributes.

**Figure 2.**
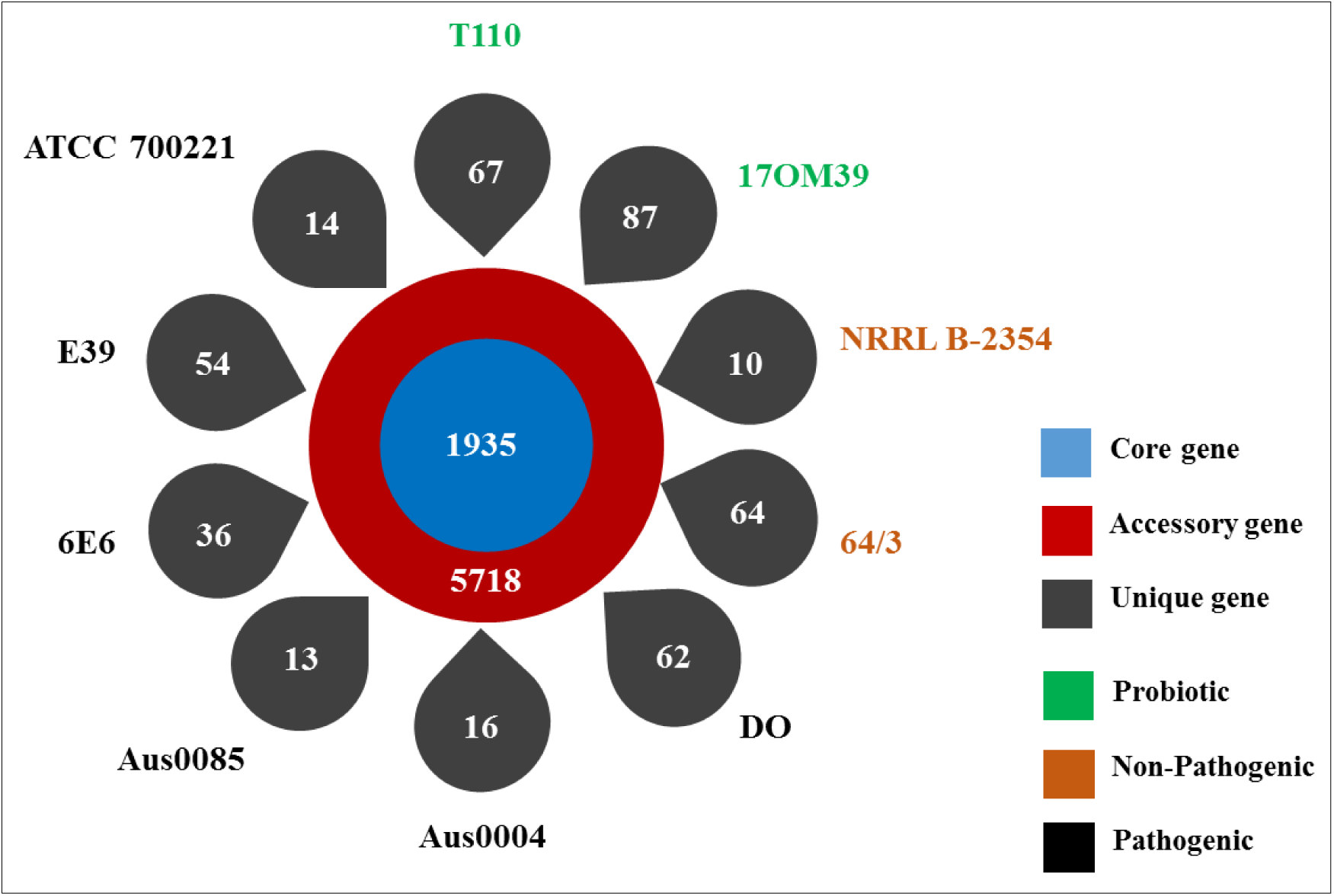
Number of Core, Accessory and Unique gene families of *Enterococcus* genomes. The inner circle represents the core genome consisting of 1935 genes in single copy. The outer red circle represents the accessory genome for all then ten strains adding to a sum of 5718 genes, while the outer petals represents the unique genes associated with all the strains. The strains green coloured are probiotics, brown are non-pathogenic and the black are pathogenic.

orthoMCL analysis of **core genes** led to the identification of 850 genes present in single copy and 772 genes present in multiple copies across all ten strains. Functional analysis of the core genes showed distribution in a varied range of functional categories within Cluster of Orthologous Genes (COG) viz. growth, DNA replication, transcription, translation, carbohydrate and amino acid metabolism, stress response and transporters. In contrast to earlier reports, core genes were also found in the functional categories of secondary metabolism and motility (Palmer et al. 2012; Beukers et al. 2017). Categories representing transport and metabolism of coenzyme, lipid, amino acid, nucleotide comprised of 16.24% of the core genes, while 11.30% of core genes were ascribed to carbohydrate metabolism which is in agreement with an earlier report (Beukers et al. 2017).

Functional analysis of the **accessory genes** showed diverse distribution in COG categories as similar to core gene annotations (Figure S3). An important observation was seen in subsystems a) carbohydrate metabolism and b) replication, recombination and repair systems. The former was abundant in the probiotic group (p = 0.002) while latter in the pathogenic group (p = 0.039) (Figure 3). This has been attributed to the properties of probiotic strains to utilize various carbohydrates(Ghattargi et al. 2018); while the pathogenic group had higher abundance of replication and recombination genes known to be associated with a large number of mobile elements(Lam et al. 2012; Qin et al. 2012). We also made an attempt to find the accessory genes being shared between the groups. The probiotic and pathogenic group shared 15 genes; four of them are general transporters, two are manganese-containing catalase gene, which provides resistance to hydrogen peroxide present in human GIT (Figure S2B) (Wang et al. 2014; King et al. 2000; Xu et al. 2016).

**Figure 3.**
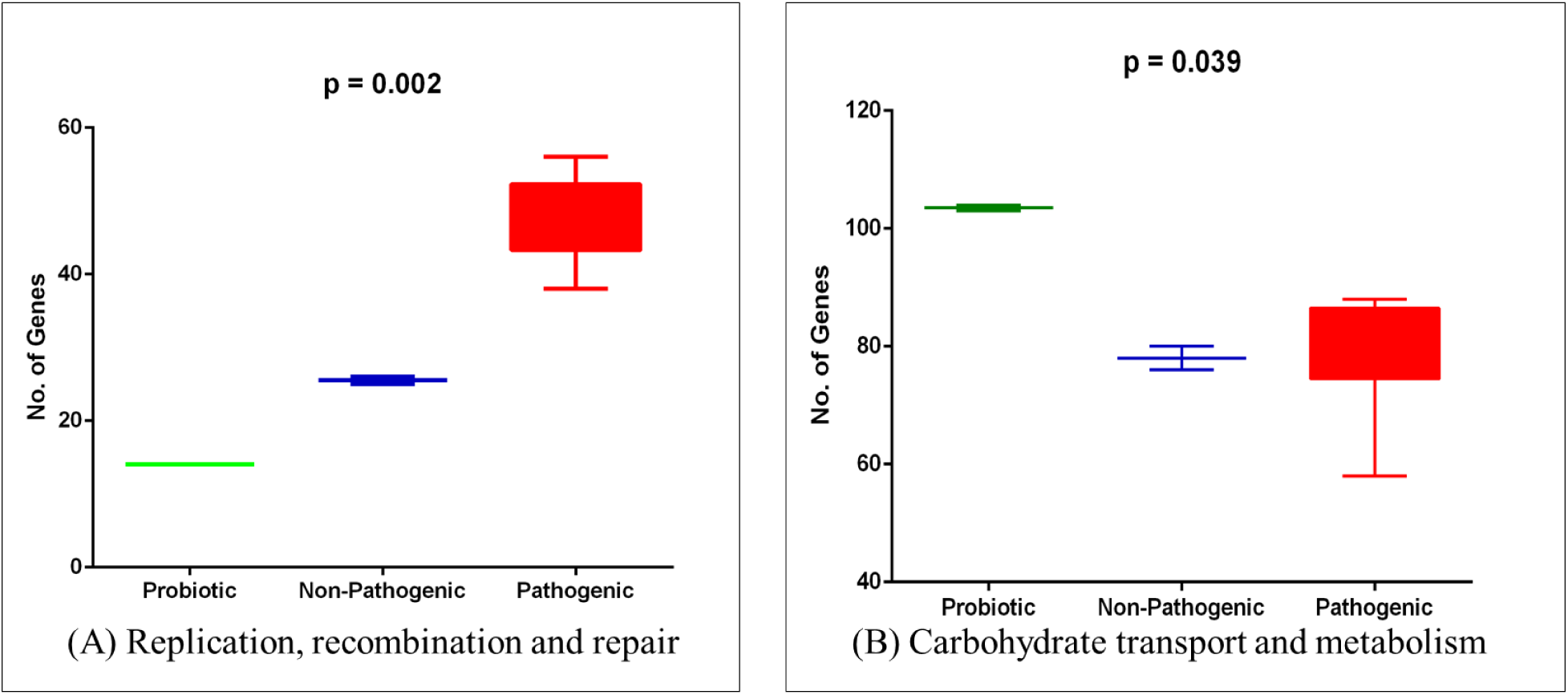
Showing the Significant COG’s in the accession genome. (A) Replication, recombination and repair (B) Carbohydrate transport and metabolism

The important **unique genes** associated with the various strains are as follows: phosphotransferase (PTS) system for mannose/fructose/sorbose in probiotic strain 17OM39 is involved in sugar uptake (Postma et al. 1993; Monot et al. 2011). Marketed probiotic strain T110 has macrolide-efflux transmembrane protein which acts as drug efflux pump and plays a key role in drug resistance (Sangvik et al. 2005; Stadler and Teuber 2002). The important unique genes for others are hexosyltransferase in strain 64/3, type III restriction-modification system in strain NRRL B-2354, Cro/CI family transcriptional regulator protein in strain 6E6, transposase for insertion sequence IS1661 in strain ATCC 700221, streptogramin A acetyltransferase in strain Aus0004, Patatin-like proteins in strain Aus0085, IS1668 transposase in strain DO, plasmid recombination enzyme in strain E39. An additional file gives the detailed information on core, accessory and unique genes in more detail [see Additional file 1, 2, 3].

Thus, the analysis conducted here has shown that pan-genome of *E. faecium* constructed on the basis of 10 genomes is still open, while the core genome seems to have reached almost a closed state. The small size of the core genome and a huge number of accessory genes support the observation of the genomic fluidity of *E. faecium* (Bakshi et al. 2016).

### Antibiotic resistance determinants

*Enterococci* can exhibit resistance to a number of antibiotics, which has been attributed to their innate resistance and also due to their ability to successfully acquire resistance through horizontal gene transfer (HGT) (Tong et al. 2017; Hegstad et al. 2010a). Multiple-drug-resistant strains of *E. faecium* have been increasingly associated with nosocomial infections particularly the vancomycin resistance (McArthur et al. 2013). Thus screening of antibiotic resistance determinants in genomes was necessary in order to understand if probiotic strains harboured these genes.

Here we screened the genomes for antibiotic resistance genes using Comprehensive Antibiotic Resistance Database (CARD) (McArthur et al. 2013). Genes conferring resistance to kanamycin were found in all the genomes which have been attributed to the intrinsic property within *E. faecium* (Galimand et al. 2011). The non-pathogenic group showed the presence of general multidrug transporter. Within pathogenic strains, Aus004 and Aus0085 showed the presence of tetracycline, trimethoprim and vancomycin resistance gene. Strains E39 and 6E6 showed the presence of genes responsible for trimethoprim and tetracycline resistance. Pathogenic strain E39 presented daptomycin resistance gene and strain ATCC 700221 showed the presence of genes responsible for resistance to the antimicrobial activity of cationic antimicrobial peptides and antibiotics such as polymyxin. Table 2 shows various antibiotic resistance genes found in each strain.

**Table 2.**
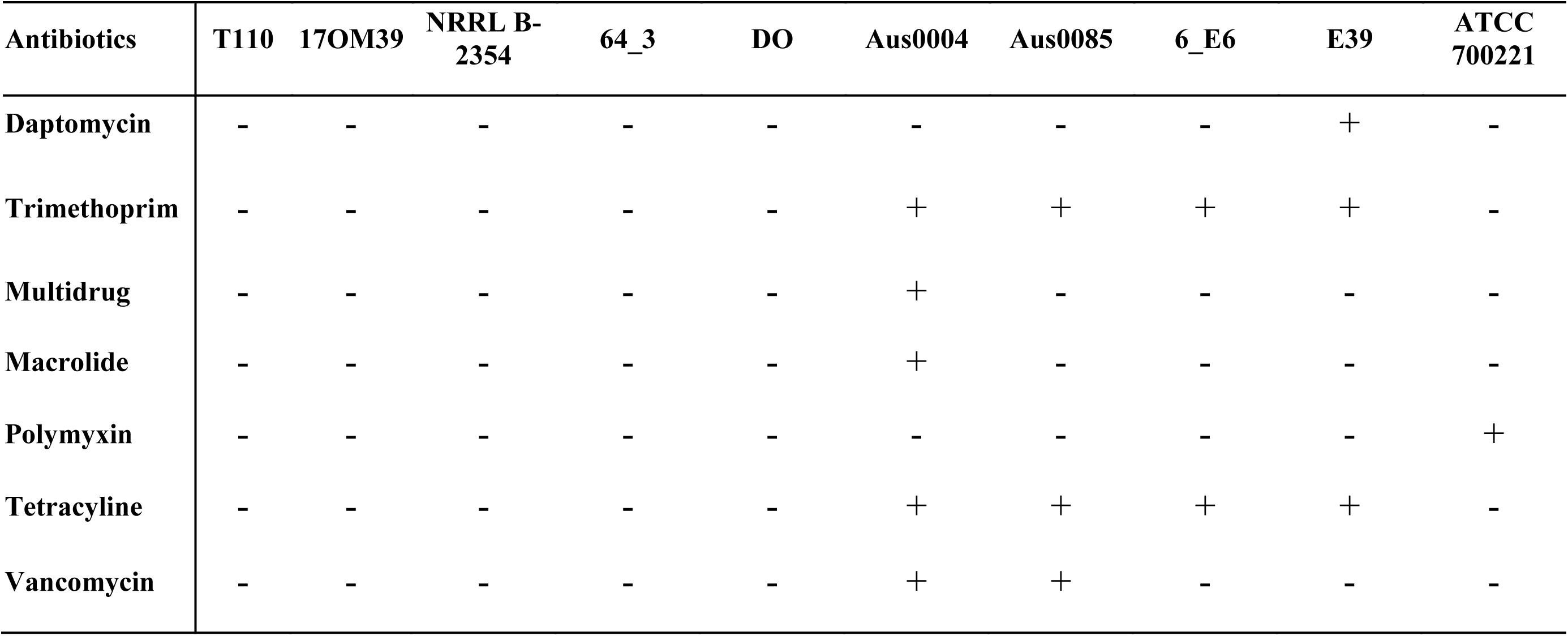
Antibiotic Resistance genes found in *Enterococcus* genomes as performed by CARD analysis, where + Present and - Absent

In our study, genes imparting resistance to one or more antibiotics were seen in different strains of *E. faecium*. Overall, the pathogenic group of *E. faecium* was found to have a higher prevalence of antibiotic resistance genes; a factor that contributes to the challenge of selecting therapeutic measures. The probiotic group was devoid of any major clinically relevant antibiotic resistance (Natarajan and Parani 2015; Ghattargi et al. 2018).

### Virulence determinants

Virulence genes contribute to the pathogenicity of an organism (Comerlato et al. 2013). Despite the increasing knowledge of *E. faecium* as an opportunistic pathogen, the distribution of virulence factors is still poorly understood (Comerlato et al. 2013). Knowledge of the virulence characteristics helps to understand the complex pathogenic process of the pathogenic strains. This study also determines genes responsible for virulence factors such as adherence, biofilm formation and exo-enzyme production in probiotic, NPNP and pathogenic groups.

The ability to adhere to the GIT is reflected to be one of the main selection criteria for potential probiotics as it extends their persistence in the intestine (Ouwehand et al. 1999) and thus allows the bacterium to exert its probiotic effects for an extended time. However, adhesion is also considered a potential virulence factor for pathogenic bacteria (Kirjavainen et al. 2001). The intestinal mucus is an important site for bacterial adhesion and colonization (Finlay 1997) and thus adherence property is beneficial to humans in case of probiotics and it possesses adverse effects in pathogenic strains. The genes described as adherence factors (*acm, scm, EbpA, EbpC*) have been attributed to pathogenic effects and our study could find most of these genes in the pathogenic group. Excluding strain DO all other pathogenic strains showed the presence of enterococcus surface protein (*esp)* gene which contributes as a major virulence factor (Heikens et al. 2007; Ramadhan and Hegedus 2005; Toledo-Arana et al. 2001; Baldassarri et al. 2001).

The *bopD* gene involved in biofilm was intact in all groups but the operon was absent in strain 17OM39, marketed probiotic strain T110 (Ghattargi et al. 2018) and non-pathogenic strains (Natarajan and Parani 2015). In an exo-enzyme group, hyaluronidase gene was found to be associated with marketed probiotic strain alone, while the gene in strain 17OM39 displayed an alteration in sequence at position 167 (G>T) suggesting this could affect its functionality due to the nonsense mutation. Gene *acm* in the probiotic and the non-pathogenic group was not functional due to the non-sense mutation at position 1060 (G>T). Also, the virulence gene *scm, efaA* and *srtC* are not well characterized as virulence determinants in *E. faecium* (Natarajan and Parani 2015) (Table 3).

**Table 3.** Virulence factors found in *Enterococcus* genomes, where + Present; - Absent; * Non-functional due to presence of stop codon.

| CATEGORY | GENES | T110 | 17OM39 | NRRL B-2354 | 64/3 | DO | Aus004 | Aus0085 | 6E6 | E39 | ATCC 700221 |
| --- | --- | --- | --- | --- | --- | --- | --- | --- | --- | --- | --- |
| Adherence | *acm* | * | * | * | * | + | + | + | + | + | + |
|  | *EbpA* | - | - | + | + | + | + | + | + | + | + |
|  | *EbpC* | - | - | + | + | + | + | + | + | + | + |
|  | *srtC* | + | + | + | + | + | + | + | + | + | + |
|  | *EcbA* | - | - | - | - | + | + | + | + | + | + |
|  | *efaA* | + | + | + | + | + | + | + | + | + | + |
|  | *Esp* | - | - | - | - | - | + | + | + | + | + |
|  | *Scm* | * | * | * | * | + | + | + | + | + | + |
|  | *SgrA* | - | - | - | - | + | + | + | + | + | + |
| Bioflim | *bopD* | * | * | * | * | + | + | + | + | + | + |
| Exoenzymes | *EF0818* | + | * | - | - | - | - | - | - | - | - |

Although one could expect a virulence trait depending on the source of isolation, our study did not find any such traits and differs from the earlier reports (Dahlén et al. 2012). Also, the strains showed significantly different patterns of virulence determinants, which underlines the findings of other author (Dahlén et al. 2012). The strain 17OM39 within the probiotic group was devoid of any clinically relevant functional virulence determinants.

### Mobile genetic elements

Mobile genetic elements (MGEs) play an important role in HGT of genes within and between bacteria (Beukers et al. 2017; Jiang et al. 2017; Kaplan 2014; Von Wintersdorff et al. 2016). A number of MGEs have been described in *E. faecium* including transposons, plasmids, and bacteriophage (Hegstad et al. 2010b; Beukers et al. 2017).

#### Insertion sequences (ISs)

are possibly the smallest and most independent transposable elements, thus playing an important role in shaping the bacterial genomes (Siguier et al. 2014). Based on the screening for IS elements (Table S1), the IS1542 was present only in probiotic strains and earlier studies on IS1542 have shown its presence in just 2 out of 65 human pathogenic strains, suggesting no direct relation with the strains pathogenicity (Huh et al. 2004). The IS element ISEfa12 was present only in non-pathogenic group and IS1216, 1S1216E, 1S1216V, IS16, IS6770, ISEf1, ISEfa10, ISEfa11, ISEfa5, ISEfa7, ISEfa8, ISEnfa3, ISS1W were present only in all the strains of the pathogenic group. The presence of insertion sequence families in all groups imply these elements are spread by HGT (Mikalsen et al. 2015). However, particular IS elements are distributed in only one group suggesting that these IS elements have evolved over the time (Werner et al. 2011; Mikalsen et al. 2015). Notably, presence of IS16 has been used as a marker within the hospital strains of *E. faecium* with 98% sensitivity and 100% specificity (Mikalsen et al. 2015). This observation was further supported by detecting IS16 in the only pathogenic group of *E. faecium* strains. Moreover, ISEfa11 and ISEfa5 are associated with vancomycin resistance genes viz. *VanS, VanX,* and *VanY* [34,65]. This correlation was also seen in this study (Figure 4).

**Figure 4.**
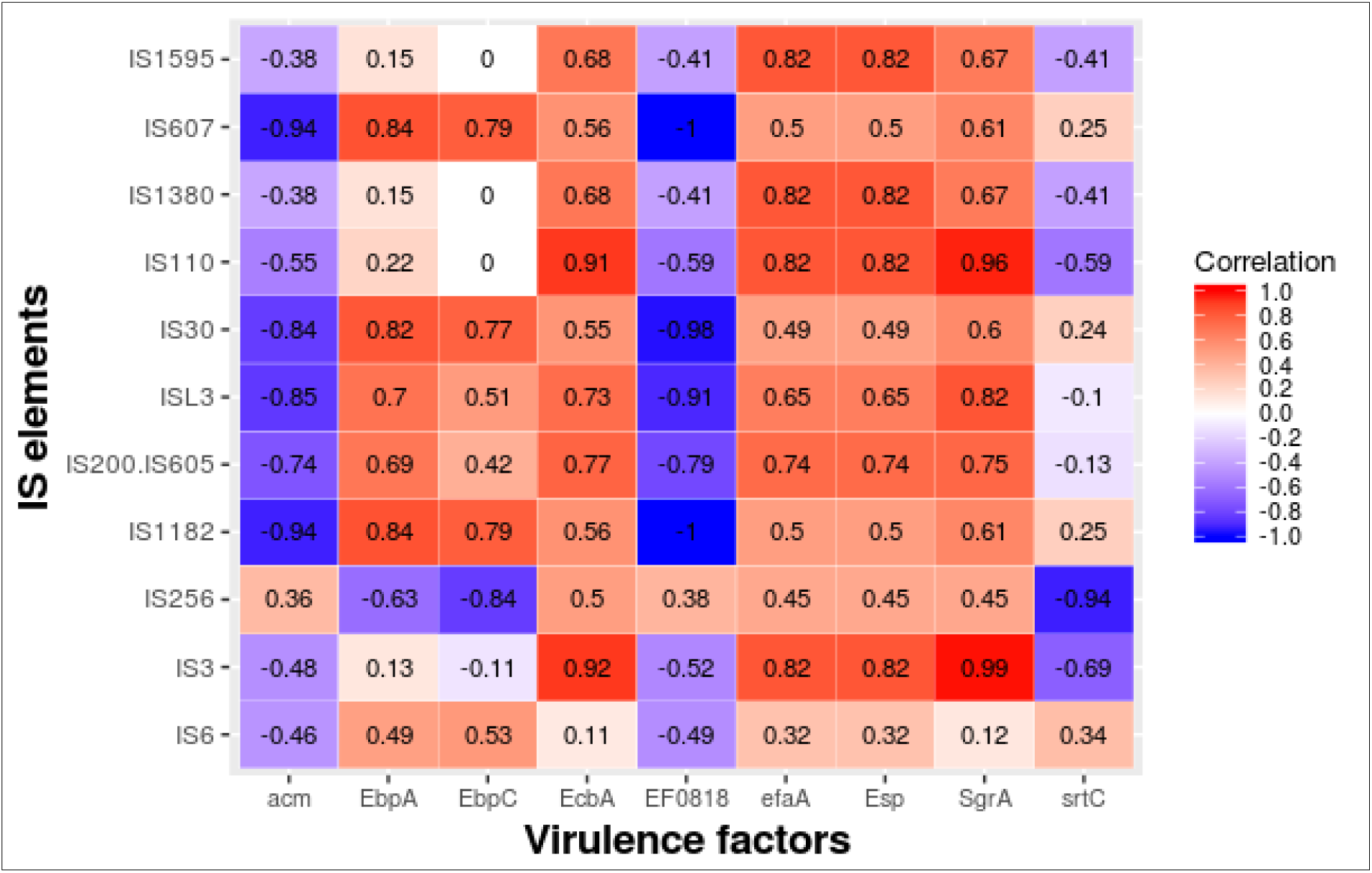
The heat map showing the correlation between the IS elements and virulence factors found across the genome. Red color indicated the strong positive correlation while the blue indicates negative correlation.

*E. faecium* are known to harbour bacteriophages, so the presence of **prophage** was predicated in all the ten genomes [34, 66]. Bacteriophages contribute actively to bacterial evolution by integrating and excising from the genome (Matos et al. 2013). In certain conditions they provide new genetic properties to the bacterial host and leading to the development of new pathogens within species, as shown for *Escherichia coli*, *Vibrio cholera* and *Corynebacterium diphtheriae* (Davis and Waldor 2003; Backman et al. 1983; FREEMAN 1951). We could trace at-least one intact prophage in all the ten genomes (Table S2). In total, there were three incomplete (PHASTER score < 70), twenty-three intact phages (PHASTER score < 70-90) and nine phages whose completeness status was doubtful (PHASTER score < 90). The non-pathogenic strains had two intact prophages, while the probiotic strain 17OM39 had one intact and a doubtful prophage while the other strain T110 had two intact prophages. *Enterococcus faecium* ATCC 700221 had the highest number of intact phages. We could not find any known functional virulence factors or genes associated with important functions within these bacteriophages regions. Further, Clustered Regularly Interspaced Short Palindromic Repeat (CRISPR) associated (Cas) system were found to be absent within the genomes as opposite to the closed neighbour, *E. faecalis* (Palmer and Gilmore 2010).

#### Genomic Islands

are distinct DNA fragments differing between closely related strains, which usually are associated with mobility (Dobrindt et al. 2004; Juhas et al. 2009). Genomic island comparison between strain 17OM39 with the other strain of *E. faecium* was done to find out the genes transferred by HGT (Table S3). We identified a total of 11 genomic island in strain 17OM39 amounting to 3.5% of total genome. The genomic island (GI1) was common across all groups except for strain 6E6. The choloylglycine hydrolase gene was found to be present in the genomic island of probiotic stain T110, pathogenic strain 6E6 and non-pathogenic strain 64/3. The choloylglycine hydrolase gene imparts resistance to bile salts and thus helps to survive within the gut environment (Jones et al. 2008; Begley et al. 2006). In this study, the pathogenic group showed a large number of IS elements, transposons and antibiotic resistance genes within the genomic island. Pathogenic strains Aus0004 and Aus0085 showed the presence of virulence factor *esp* gene (enterococcal surface protein) and vancomycin resistance genes and only pathogenic strains alone showed the presence of tetracycline resistance gene within the genomic island. They also showed presence of a cell adhesion protein within the Genomic Island. A higher similarity was observed between genomic islands of probiotic and NPNP strains as compared to pathogenic group (Figure 5). The distribution of these MGE‟s within the genome is as shown in Figure 6 with strain AUS0004 having nearly 25% of its genome as mobile.

**Figure 5.**
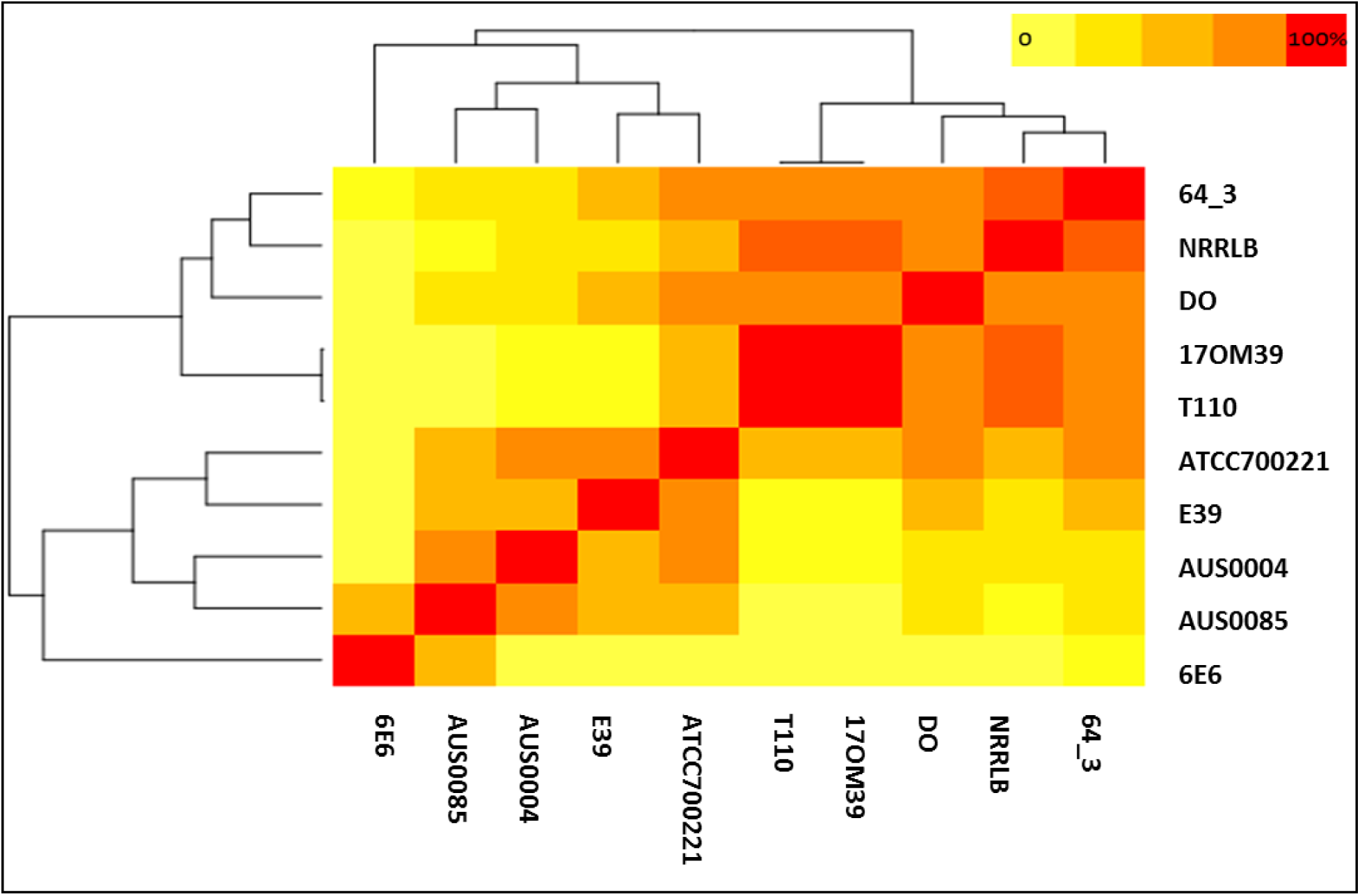
Heat map showing the similarities between the Genomic Islands of the strains considered in this study. The lightyellow shows least percent similarity while the red indicate 100% similarity with the genomes of the strains considered in the study. The pathogenic strains (E39, Aus0004, Aus0085 and 6E6) shows clustering while the probiotic strains (T110 & 17OM39), nonpathogenic strains (64_3 & NRRLB) and pathogenic strain DO present another clustering. The color scheme is as shown in the top right corner.

**Figure 6.**
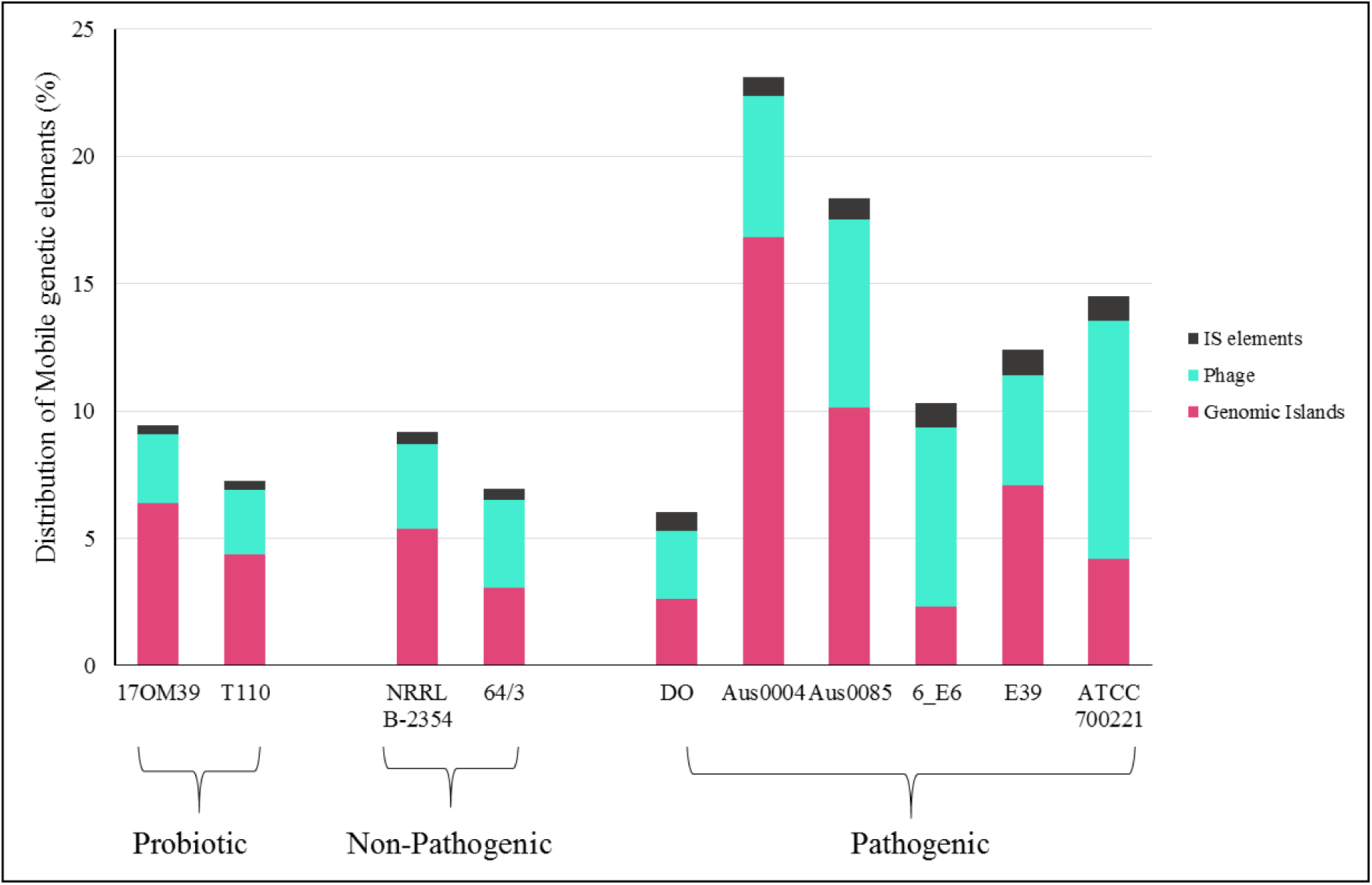
Proportion of Mobile Genetic Elements across *Enterococcus* genomes. The pick colour shows the proportion of genomic islands present in the each strain, light green for bacteriophages and black of IS elements across all the strains. Strain AUS004 has nearly quarter of its genome packed with these mobile genetic elements.

*E. faecium* behaves as probiotic, non-pathogenic and pathogenic these underlying mechanisms may be intrinsic to the strain or acquired by horizontal exchange of genetic material. Thus genes associated with Genomic Islands can be considered as acquired properties (Juhas et al. 2009; Hollenbeck and Rice 2012; Dobrindt et al. 2004; Gold 2001; Marothi et al. 2005). Genes associated with probiotic properties in strain 17OM39 were found to be associated with the genome and not within any Genome islands. From this study, it is evident that the role of the MGEs in adding new capacities and also driving the evolution of the strains (Jones et al. 2008; Begley et al. 2006). Certainly, studies like one carried out here are helpful to understand the evolution of predominant strains. An additional file gives the detailed information on genes present in Genomic Islands in more detail [see Additional file 4].

### Survival in Gastrointestinal Tract

Biologically active microorganisms are usually required at the target site to induce a health benefit or pathogenic effect, but must survive the host‟s natural barriers such as the acid, bile and must adhere to GIT (İspirli et al. 2015; Banwo et al. 2013; Ahmadova et al. 2013; Rao et al. 2013). Moreover, *E. faecium* is a part of the core microbiome and their number in human faeces ranges from 10^4^ to 10^5^ per gram (Santagati et al. 2012). Thus a list of genes encoding for survival and growth were observed in the strain 17OM39 and compared with other genomes. We found Permease IIC component gene responsible for catalysing the phosphorylation of incoming sugar substrates (Minelli and Benini 2008) only in the probiotic group. All the groups showed the presence of the genes that impart resistance to acid, bile and can hydrolyze bile salt. Moreover, these strains were also able to adhere and grow in the GI conditions. This finding correlates with the fact that *Enterococcus faecium* are normal inhabitants of the gut (Campos et al. 2004) (Table 4).

**Table 4.** Number of genes responsible for survival in GI track within *Enterococcus* genomes.

| CATEGORY | GENES | T110 | 17OM39 | NRRL B-2354 | 64/3 | DO | Aus0004 | Aus0085 | 6_E6 | E39 | ATCC 700221 |
| --- | --- | --- | --- | --- | --- | --- | --- | --- | --- | --- | --- |
| Acid resistance | *LBA0995*<br>*LBA1524*<br>*LBA1272*<br>*gadC*<br>*rrp-1* | 3/5 | 3/5 | 3/5 | 3/5 | 4/5 | 4/5 | 5/5 | 4/5 | 4/5 | 3/5 |
| Bile resistance | *LBA1430*<br>*clpE*<br>*dps*<br>*LBA1429* | 4/4 | 4/4 | 4/4 | 4/4 | 4/4 | 4/4 | 4/4 | 4/4 | 4/4 | 4/4 |
| Competitive | *copA*<br>*met*<br>*pts14C* | 3/3 | 3/3 | 2/3 | 2/3 | 2/3 | 2/3 | 2/3 | 2/3 | 2/3 | 2/3 |
| Adherence | *lsp*<br>*FbpA*<br>*ispA* | 2/3 | 2/3 | 3/3 | 3/3 | 3/3 | 3/3 | 3/3 | 3/3 | 3/3 | 3/3 |
| Persistence | *LJ1656*<br>*msrB*<br>*LJ1654*<br>*clpC* | 4/4 | 4/4 | 4/4 | 4/4 | 4/4 | 4/4 | 4/4 | 4/4 | 4/4 | 4/4 |
| Bile salt hydrolase | *bsh* | 1/1 | 1/1 | 1/1 | 1/1 | 1/1 | 1/1 | 1/1 | 1/1 | 1/1 | 1/1 |
| Growth | *treC* | 1/1 | 1/1 | 1/1 | 1/1 | 1/1 | 1/1 | 1/1 | 1/1 | 1/1 | 1/1 |
| Adaptation | *Lr1265*<br>*Lr1584* | 1/2 | 1/2 | 1/2 | 1/2 | 1/2 | 2/2 | 2/2 | 2/2 | 2/2 | 2/2 |

### Probiotic properties

Numerous characteristics should be taken into account when selecting a probiotic strain. Some examples of desirable characteristics for a probiotic strain include the ability to survive and retain viability in acid and bile concentrations, simulating the harsh environment of GIT (Charteris et al. 1998; Rehaiem et al. 2014). A probiotic strain should have the ability for producing antimicrobial substances but be devoid of acquired antibiotic resistance genes (Charteris et al. 1998; Rehaiem et al. 2014; Fao et al. 2002; Ganguly et al. 2011). As stated earlier the strain 17OM39 and marketed probiotic strains T110 were devoid of any clinically relevant antibiotic resistance gene while all the strains were able to survive in GIT conditions.

For strains 17OM39 and T110 (marketed probiotic) we could trace complete pathways for amino acid synthesis viz. valine, lysine, and methionine (Table 5). These are among the essential amino acids and need to be supplied exogenously (Food and Nutrition Board 1989). Vitamins such as folate and thiamine are the components of Vitamin-B and are considered as essential nutrients for humans. Folate (folic acid) cannot be synthesized by human cells and hence is necessary to be supplemented exogenously as it plays important role in DNA and RNA synthesis and amino acid metabolism (Mahmood 2014; Ohrvik and Witthoft 2011; Tuszyńska 2012). Thus, strains (T110 and 17OM39) producing such amino acids and vitamins are considered beneficial for human use (Fijan 2014; Eck and Friel 2013). Genes responsible for antibacterial activity (bacteriocin) specific against *Listeria* were found. Genes for exopolisaccharide (EPS) and anti-oxidant (hydro-peroxidases) production were noted which in-turn help the probiotic strains in establishing themselves in the gut. The non-pathogenic group only had EPS gene cluster. Complete pathways for amino acid and vitamin synthesis were absent in NPNP and pathogenic group. Thus certain strains contribute beneficially to health as observed by the probiotic group strains.

**Table 5.** Probiotic properties found in *Enterococcus* genomes, where + Present and - Absent

| Properties | 17OM39 | T110 | 64/3 | NRRL<br>B-2354 | DO | Aus0004 | Aus0085 | 6_E6 | E39 | ATCC<br>700221 |
| --- | --- | --- | --- | --- | --- | --- | --- | --- | --- | --- |
| Anti-oxidant | + | + | - | - | - | - | - | - | - | - |
| Anti-bacterial | + | + | - | - | - | - | - | - | - | - |
| EPS | + | + | + | + | - | - | - | - | - | - |
| Amino-acid | valine, lysine,<br>methionine | valine,<br>lysine | - | - | - | - | - | - | - | - |
| Vitamins | Folate,<br>Thiamine | Folate,<br>Thiamine | - | - | - | - | - | - | - | - |

### Plasmids

Earlier studies within *E. faecium* isolates have shown the abundance of plasmids by finding 1-7 number of plasmids in 88 out of 93 isolates (Rosvoll et al. 2010). Plasmids comprise a substantial portion of the accessory genome and are accountable for much of antibiotic and virulence traits to be acquired by the HGT (Rosvoll et al. 2010). Thus an attempt was made to compare the plasmids of the strains considered in this study with respect to their virulence factors, antibiotic resistance, phage regions and IS elements.

Of all the strains taken for this study, we could find a gene with 66% similar to the cytolysin (*cyl*) gene in probiotic strain T110. Studies with the *cyl* gene have been an important determinant in lethality of endocarditis (Rosvoll et al. 2010). The CARD analysis of the plasmids shows the resistance to antibiotics viz vancomycin, streptothricin, erythromycin, gentamicin and kanamycin by pathogenic group stains: 6E6, ATCC700221, Aus0085, and E39 only (Table S4). No phage elements were associated with the probiotic strain T110, while the non-pathogenic strain (NRRL B-2354) and pathogenic strains (ATCC 700221, Aus0085, DO, and E39) harboured incomplete or complete prophage. The list of IS elements found in the plasmids of strains is summarised in the TableS5.

### Delineating Probiotic, Non-pathogenic and Pathogenic strains

Multi-Locus Sequence Analysis (MLSA) based phylogeny could not distinguish between pathogenic and non-pathogenic strains of *E. faecium* (Homan et al. 2002), but this could be achieved on the basis of the core genome SNP based phylogeny (Sankarasubramanian et al. 2016; Heydari et al. 2013; Chen et al. 2013; Vliet and Kusters 2015). Thus, concatenated 1935 core genes were used to construct a phylogenetic tree of the 10 strains along with *Enterococcus faecalis* symbioflor as an out-group. Phylogenetic reconstruction by using Maximum likelihood method separated 10 strains in 3 distinct clusters as the three groups considered (bootstrap >90) (Figure 7). We found no clustering based on the source of isolation, while strain 17OM39 was closely related to the probiotic strain T110. The same observation was made when repeated with the pan-genome (data not shown).

**Figure 7.**
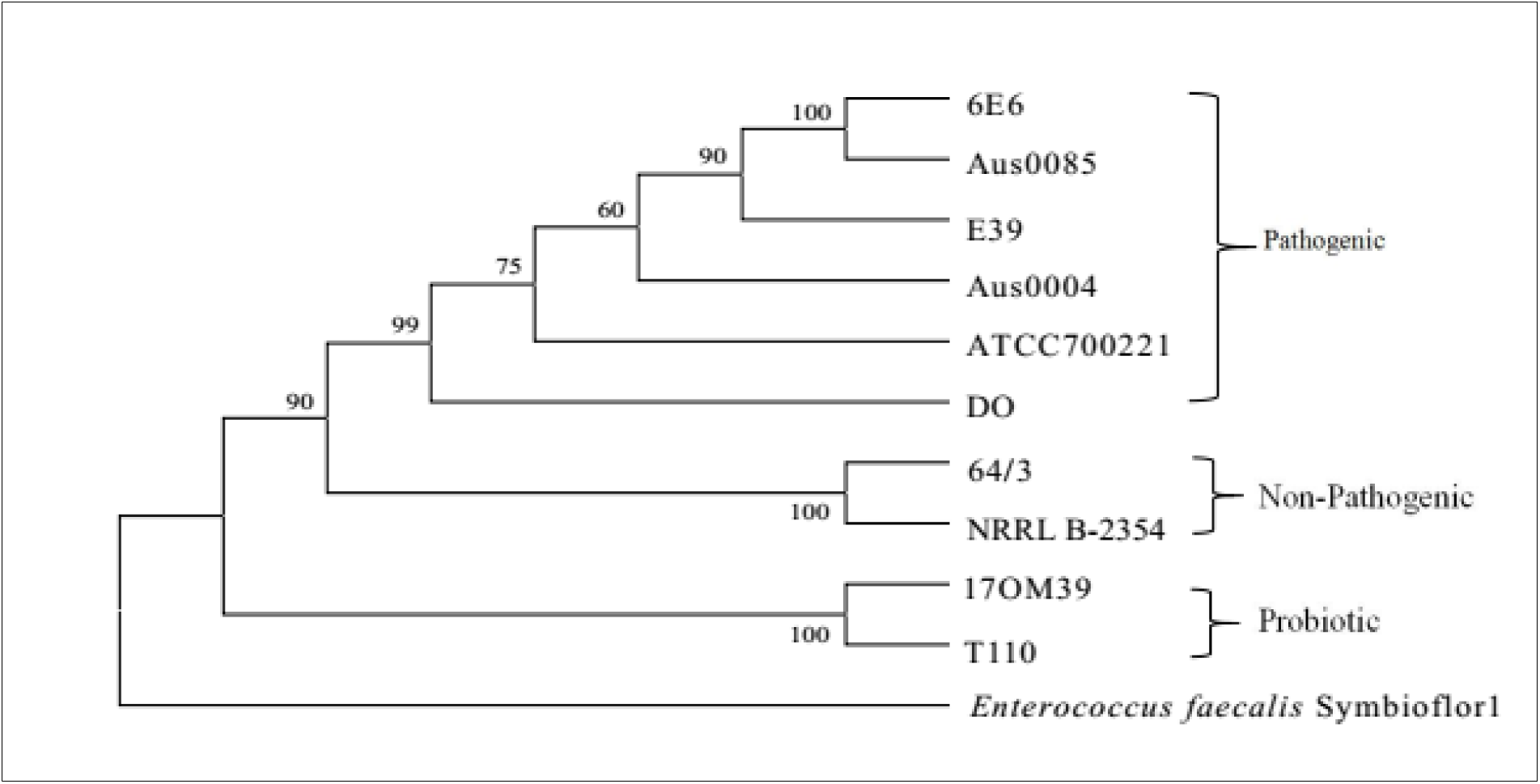
Core Genome Phylogeny. Phylogenetic tree of 10 *Enterococcus faecium* strains using the Maximum Likelihood method based on the GTR + G substitution model. The tree with the highest log likelihood (-17644.1414) is shown. Evolutionary analyses were conducted in MEGA6. A concatenated tree of 1945 genes was considered in the final dataset.

PCA plot was generated using Euclidean distances based on the prevalence of genes in antimicrobial resistance, survival in GIT and probiotic properties. The PCA plot representing the differences in property harboured between the groups is as shown in Figure 8. The BLAST Atlas was generated with the help of GVIEW server with strain Aus0004 as the reference genome (Figure 9). Of the strains, Aus0085 exhibited the highest relatedness to the reference strain. There were variable regions identified among non-pathogenic, pathogenic and probiotic group illustrating their dissimilarity in genome content. Pathogenic Island (2812458-2878042 and 1860143-1894650 bp) consist of majorly virulence-associated genes, IS elements transposes and integrase and antibiotic resistance-related genes. It also has vancomycin resistance gene cluster and presence of *esp* gene which correlates with the previous studies (Shankar et al. 2002; McBride et al. 2009). However, several phages and transposon-related loci from the reference strain appeared to be absent in marketed probiotic T110 and 17OM39 strains. This observation further supports their distinct segregation into independent clades.

**Figure 8.**
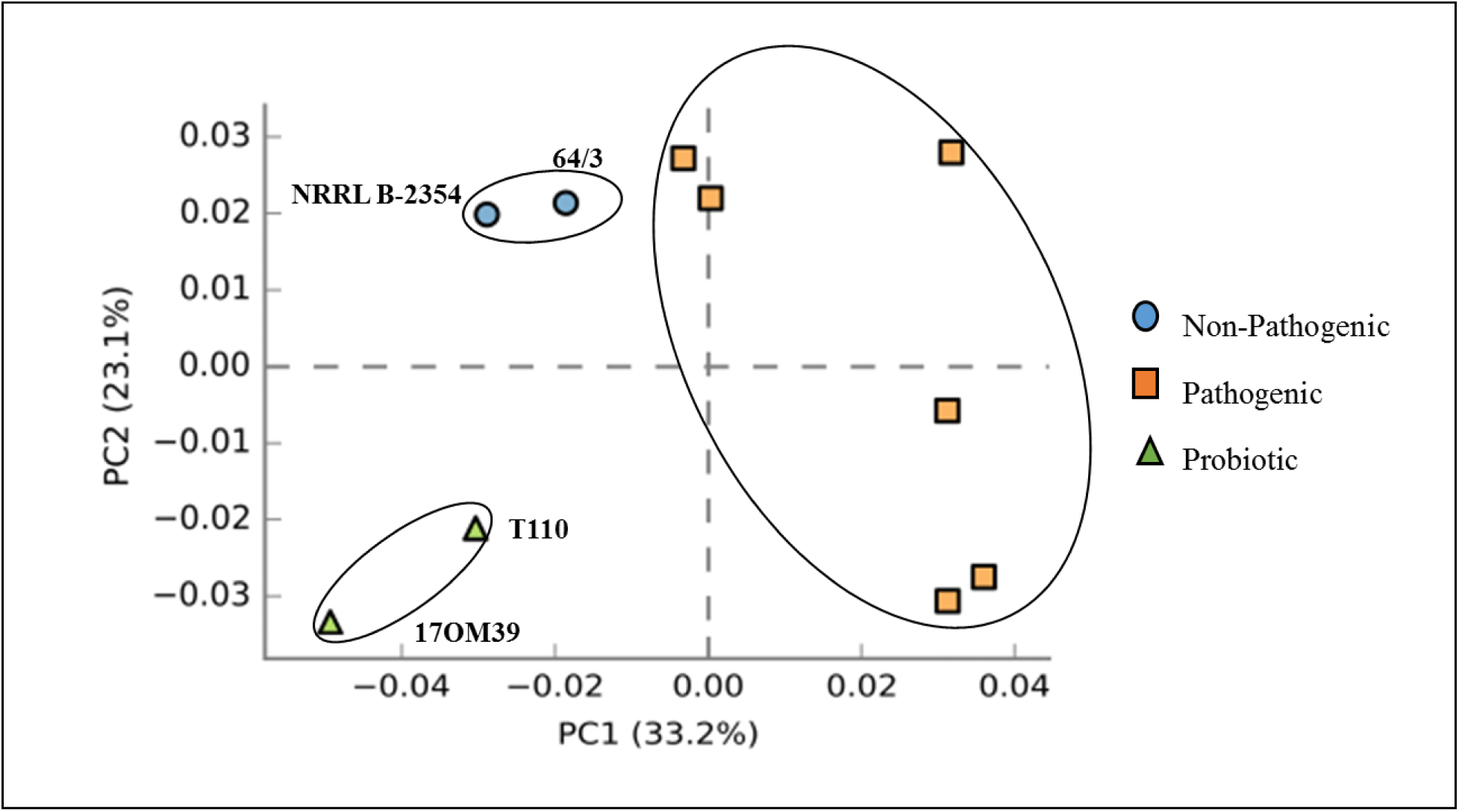
PCA plot comparing probiotic, pathogenic and non-pathogenic *Enterococcus* genomes based on presence and absence of genes responsible for survival in GI track, virulence factors and antibiotic resistance. The probiotic strains are shown in green, non-pathogenic in blue and pathogenic in red colour and the clustering is indicated by the oval shaped rings on the strains. From the plot, it can be noted that strain 17OM39 is different from the marketed probiotic strain T110.

**Figure 9.**
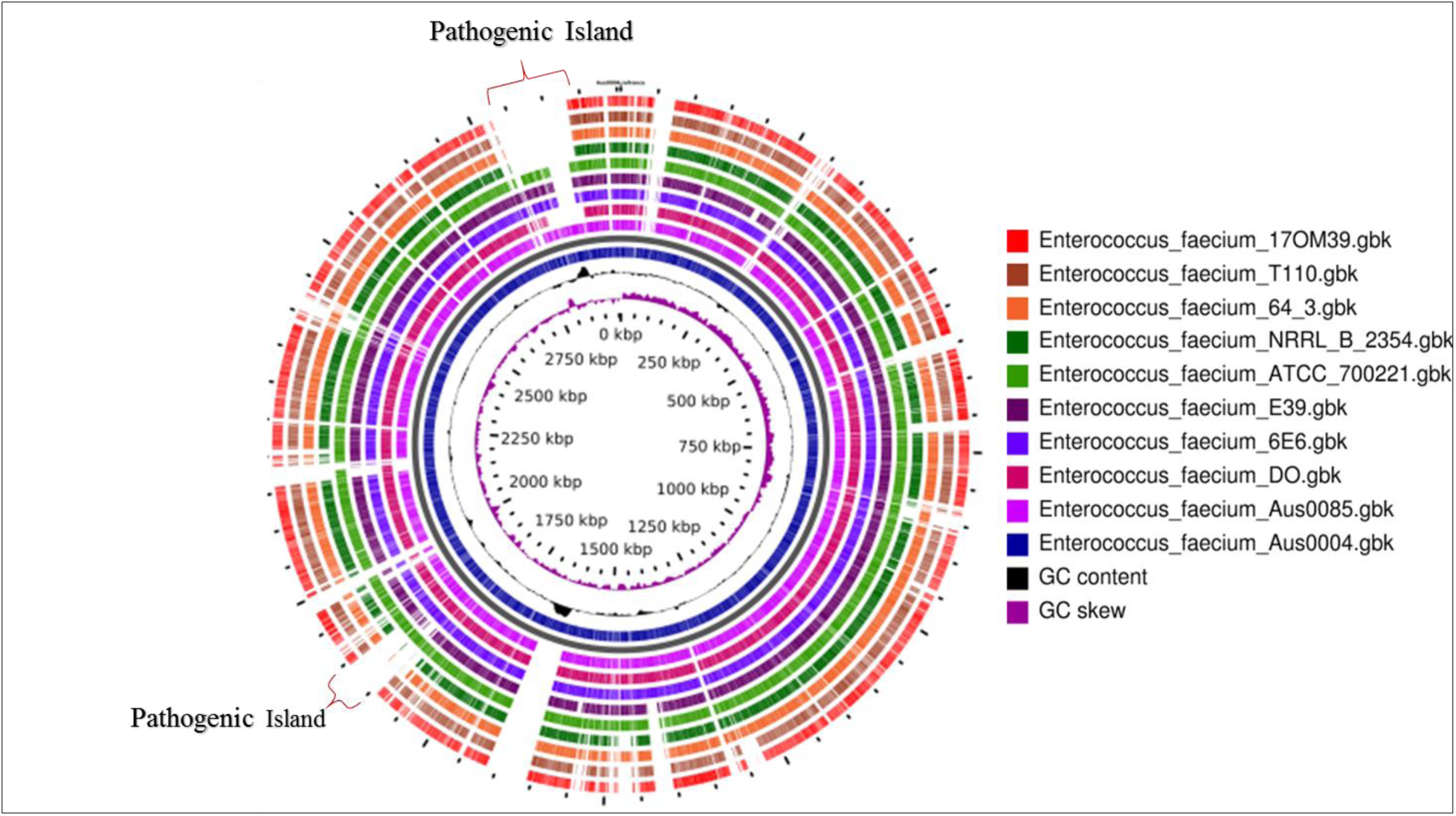
Blast Atlas of *Enterococcus* genomes, with strain Aus004 as a reference genome followed by Aus0085, DO, 6E_6, E_39, ATCC_7200221, NRRLB_2354, 64_3, T110 and the outermost as 17OM39. The two pathogenic islands (has most of virulence factors and antibiotic resistance genes) are shown in figure.

## CONCLUSIONS

This study has provided a valuable insight into genome-based investigations and has refined our knowledge on the genomic diversity of probiotic, non-pathogenic and pathogenic strains of *E. faecium*. The analysis has helped us to define core and accessory genome and also to understand the genomic relationships within them. The genomic features responsible for survival in GI tract, antibiotic resistance, virulence factors were known. We also have highlighted an abundance of mobile elements including prophage, insertion sequence elements, genomic islands, and plasmids. Moreover, the analysis of intrinsic and acquired properties helped us to know the inherent probiotic properties of strain 17OM39.

## MATERIAL AND METHODS

### Bacterial sequences and strains

Whole Genome Sequence of *E. faecium* was retrieved from NCBI genomes and a total of ten strains were used in this study. All the genomes were RAST annotated (Overbeek et al. 2014).

### Comparative analysis

Comparative analysis of ten whole genome sequences of *Enterococcus faecium* was done by an ultra-fast bacterial pan-genome analysis pipeline (BPGA) (Chaudhari et al. 2016) which performs GC content analysis, pan-genome profile analysis along with sequence extraction and phylogenetic analysis. Furthermore, the genome was investigated for the presence of putative virulence genes using Virulence Factor of Bacterial Pathogens Database (VFDB)(Chen et al. 2005). Screening of probiotic genes was done by performing BLAST of probiotic genes to the genome by online NCBI‟s BLASTX tool (Altschul et al. 1990). Comprehensive Antibiotic resistance Database (CARD) was used for analysis of antibiotic resistance (McArthur et al. 2013). Presence of CRISPR repeats was predicted using the CRISPRFinder tools (Grissa et al. 2008). PHASTER: a rapid identification and annotation of prophage sequences within bacterial genomes were used for identification of prophages within the genome (Arndt et al. 2016). Bacterial insertion elements (ISs) were identified by ISfinder (Siguier et al. 2006). Horizontal gene transfer was detected by genomic island tool: Islandviewer (Langille and Brinkman 2009; Dhillon et al. 2015). The clustering and annotation of protein sequences were done with the help of orthoMCL (Li et al. 2003). COG analysis was done with the help of webMGA server (Wu et al. 2011). STAMP software was used to generate PCA plot (Parks et al. 2014). A blast atlas was generated with the help of GVIEW Server (https://server.gview.ca/) (Petkau et al. 2010).

## List of abbreviations

GIT: Human gastrointestinal tract
CDS: coding DNA sequence
HGT: horizontal gene transfer
CARD: Comprehensive Antibiotic Resistance Database
MGEs: Mobile genetic elements
ISs: Insertion sequences
GI: genomic island
PTS: phosphotransferase system
EPS: exopolysaccharide
BPGA: ultra-fast bacterial pan-genome analysis pipeline
VFDB: Virulence Factor of Bacterial Pathogens Database

## DECLARATIONS

### Consent for publication

Not Applicable

### Ethics approval and consent to participate

Not Applicable

### Availability of data and material

Not Applicable

### Authors’ contributions

VG, DD, SP: Designed the concept; VG, MG, DD, YM: Execution of the concept; DD, SP: Guide to the concept, BM, OP, YS, YM: checking and intellectual inputs to the paper.

## Acknowledgements

Additional financial support was provided by the Department of Biotechnology, Basaveshwar Engineering College, Bagalkot, under TEQUIP Grant to Bharati S. Meti.

## SUPPLEMENTARY MATERIAL FIGURES

**Fig S1.**
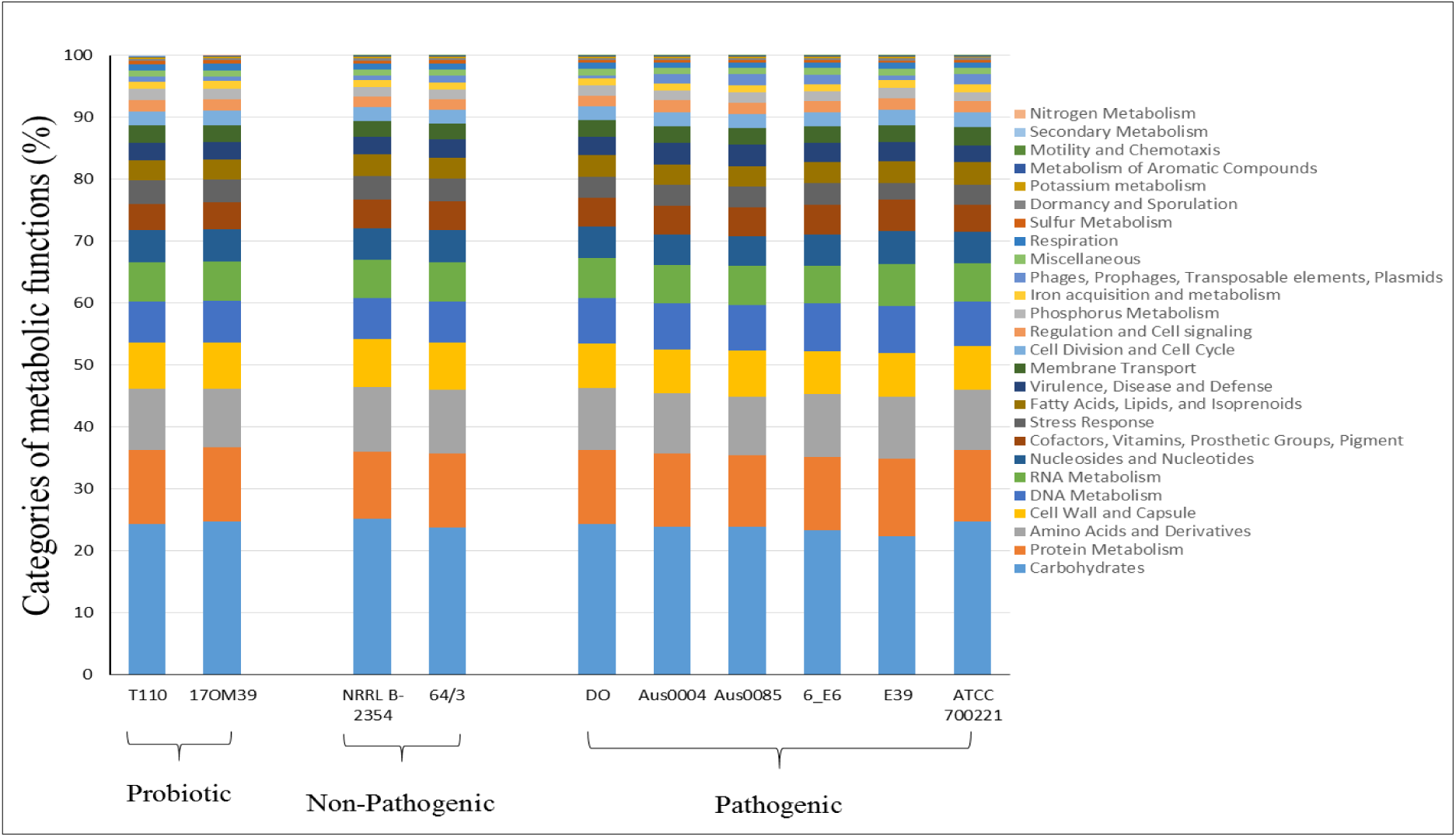
Features assigned to subsystems from RAST present in all ten *Enterococcus* strains.

**Figure S2.**
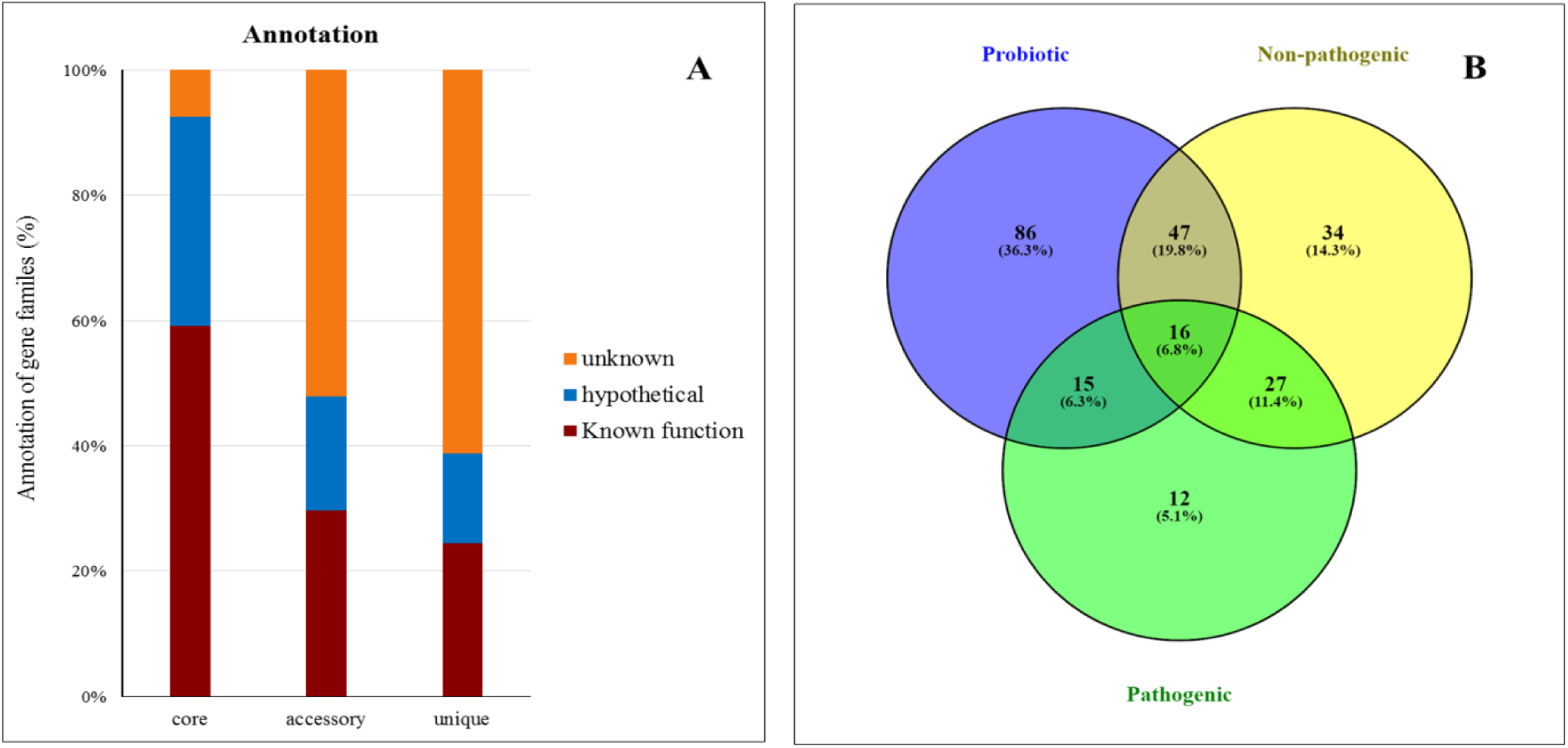
**(A)** Proportion of known, hypothetical and unknown proteins in the group of core, accessory and unique genes **(B)** Venn Diagram for accessory genome between probiotic, non-pathogenic and pathogenic group.

**Figure S3.**
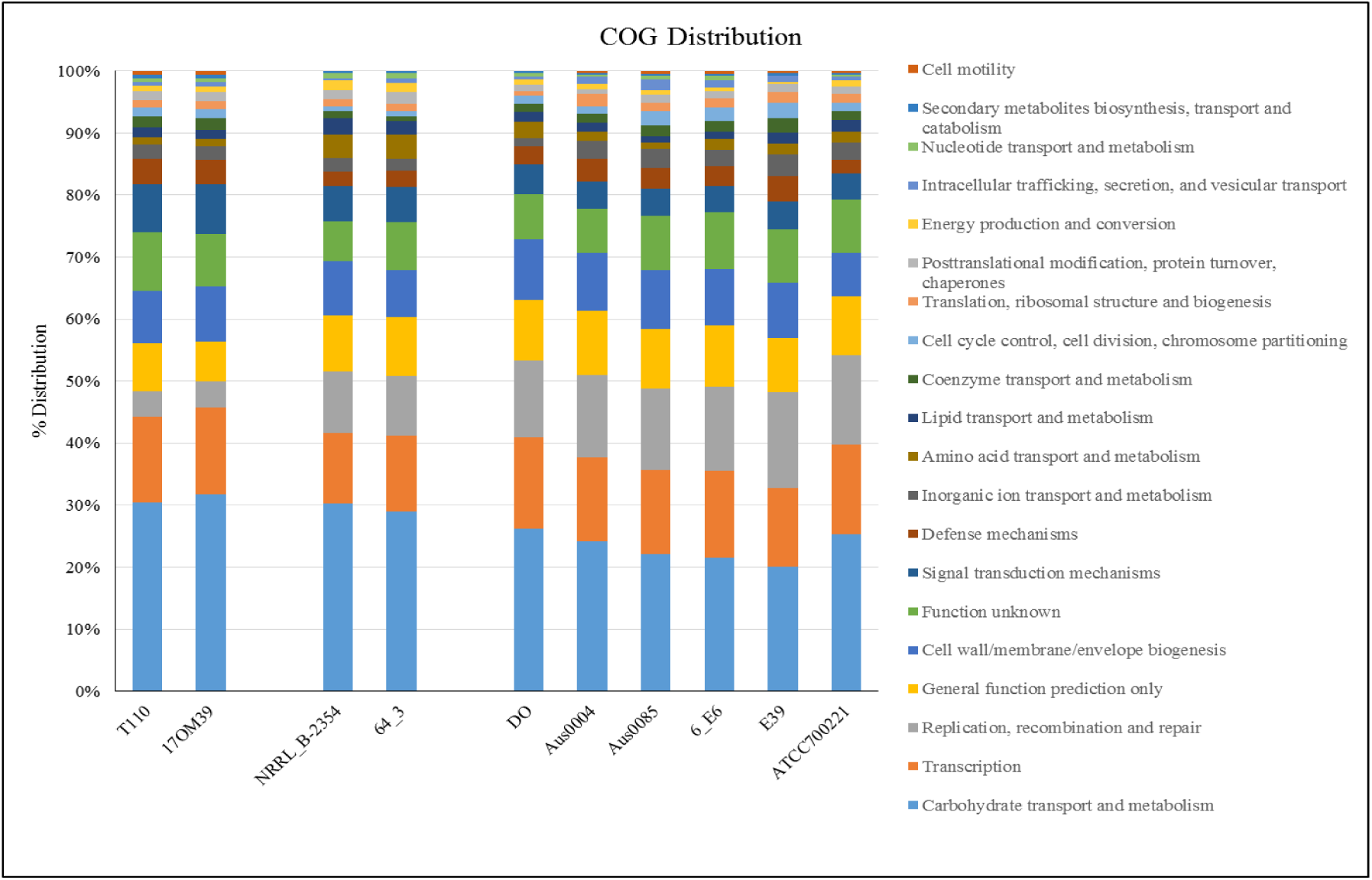
Functional analysis of the accessory genes in COG categories

**Table S1.** IS elements found in *Enterococcus* genomes by ISfinder tool. + Present, - Absent

| IS elements | Probiotic | Non-Pathogenic | Pathogenic |
| --- | --- | --- | --- |
| IS1216 | - | + | + |
| IS1216E | - | + | + |
| IS1216V | - | + | + |
| IS1251 | - | + | + |
| IS1476 | - | - | + |
| IS1485 | + | + | + |
| IS1542 | + | - | - |
| IS16 | - | - | + |
| IS1678 | - | - | + |
| IS19 | + | + | + |
| IS256 | - | - | + |
| IS6770 | - | + | + |
| ISEf1 | - | - | + |
| ISEfa10 | - | + | + |
| ISEfa11 | - | - | + |
| ISEfa12 | - | + | - |
| ISEfa13 | - | - | + |
| ISEfa4 | - | - | + |
| ISEfa5 | - | - | + |
| ISEfa7 | - | - | + |
| ISEfa8 | - | - | + |
| ISEfm1 | + | + | + |
| ISEfm2 | + | - | + |
| ISEnfa110 | - | - | + |
| ISEnfa3 | - | - | + |
| ISEnfa4 | + | - | + |
| ISLgar5 | + | - | + |
| ISS1W | - | + | + |
| ISSsu5 | - | - | + |

**Table S2.** Number of Phage elements present in *Enterococcus* genomes as intact, questionable and incomplete.

| Phage | NRRL |  |  |  |  |  |  |  | ATCC |  |
| --- | --- | --- | --- | --- | --- | --- | --- | --- | --- | --- |
|  | 17OM39 | T110 | B-2354 | 64/3 | DO | Aus0004 | Aus0085 | 6_E6 | E39 | 700221 |
| Intact | 1 | 2 | 2 | 2 | 1 | 3 | 3 | 3 | 2 | 4 |
| Questionable | 1 | - | - | - | 1 | 1 | 1 | 1 | 2 | 2 |
| Incomplete | - | - | - | - | 1 | 1 | 1 | - | - | - |
| Total Size (bp) | 73100 | 68700 | 87000 | 89100 | 74500 | 164900 | 220600 | 208400 | 117000 | 268300 |

**Table S3.** Number of Genomic Islands in *Enterococcus* genomes

| Genome | Genomic Islands | Size (bp) |
| --- | --- | --- |
| 17OM39 | 11 | 172279 |
| T110 | 9 | 117774 |
| NRRL B-2354 | 12 | 142123 |
| 64/3 | 7 | 78409 |
| DO | 7 | 73539 |
| Aus0004 | 35 | 496552 |
| Aus0085 | 24 | 304164 |
| 6_E6 | 8 | 69233 |
| E39 | 17 | 191228 |
| ATCC 700221 | 10 | 119456 |

**Table S4.** Antibiotic Resistance genes found in *Enterococcus* plasmids as performed by CARD analysis, where + Present and - Absent

| Genes | 6_E6 | ATCC_700221 | Aus0085 | DO | E39 |
| --- | --- | --- | --- | --- | --- |
| **Aminoglycoside** | + | + | + | + | + |
| **Chloramphenicol** | - | - | - | + | - |
| **Dihydrofolate** | - | + | - | - | - |
| **Erythromycin** | + | + | + | + | + |
| **Gentamicin** | + | + | - | - | - |
| **Lincosamide** | - | - | + | - | - |
| **Streptothricin** | - | + | - | + | + |
| **Tetracycline** | - | - | - | + | - |
| **Vancomycin** | + | + | - | - | + |

**Table S5.**
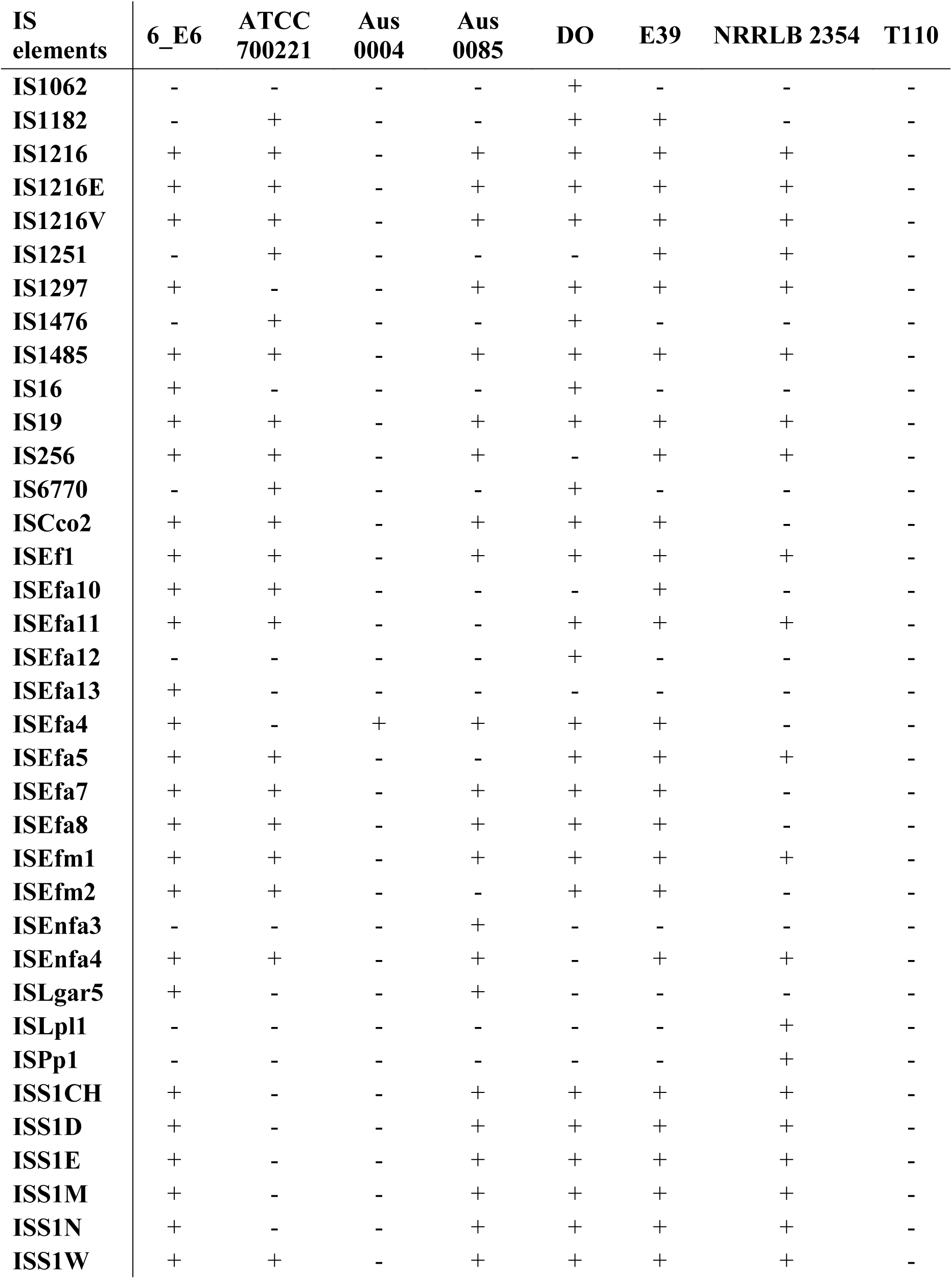
IS elements found in *Enterococcus* plasmids by ISfinder tool. + Present, - Absent

